# Coupled structural transitions enable highly cooperative regulation of human CTPS2 filaments

**DOI:** 10.1101/770594

**Authors:** Eric M. Lynch, Justin M. Kollman

## Abstract

Many enzymes assemble into defined oligomers, providing a mechanism for cooperatively regulating enzyme activity. Recent studies in tissues, cells, and *in vitro* have described a mode of regulation in which enzyme activity is modulated by polymerization into large-scale filaments^1–5^. Enzyme polymerization is often driven by binding to substrates, products, or allosteric regulators, and tunes enzyme activity by locking the enzyme in high or low activity states^1–5^. Here, we describe a unique, ultrasensitive form of polymerization-based regulation employed by human CTP synthase 2 (CTPS2). High-resolution cryoEM structures of active and inhibited CTPS2 filaments reveal the molecular basis of this regulation. Rather than selectively stabilizing a single conformational state, CTPS2 filaments dynamically switch between active and inactive filament forms in response to changes in substrate and product levels. Linking the conformational state of many CTPS2 subunits in a filament results in highly cooperative regulation, greatly exceeding the limits of cooperativity for the CTPS2 tetramer alone. The structures also reveal a link between conformational state and control of ammonia channeling between the enzyme’s two active sites. This filament-based mechanism of enhanced cooperativity demonstrates how the widespread phenomenon of enzyme polymerization can be adapted to achieve different regulatory outcomes.

---

CTP synthase (CTPS) is the key regulatory enzyme in pyrimidine biosynthesis, with critical roles in regulation of nucleotide balance^6^, maintenance of genome integrity^7,8^, and synthesis of membrane phospholipids^9^. CTPS catalyzes the conversion of UTP to CTP in an ATP-dependent process, the rate-limiting step in CTP synthesis. CTPS is regulated through feedback inhibition by CTP binding, and is allosterically regulated by GTP, making it sensitive to levels of the four essential ribonucleotides, reflecting its role as a critical regulatory node in nucleotide metabolism^10–13^. CTPS is a homotetramer, with each monomer composed of a glutaminase and an amidoligase domain connected by a helical linker^14^. Ammonia is generated from glutamine then transfered to the amidoligase domain, where it is ligated to UTP to form CTP; while both of these catalytic mechanisms are well understood, the mechanism of ammonia transfer between the two separated active sites has not yet been described. Previously, we showed that CTPS undergoes a conserved conformation cycle controlled by substrate and product binding, involving two major structural changes: upon substrate binding, the glutaminase domain rotates towards the amidoligase domain, bringing the two active sites closer, and the tetramer interface rearranges to accommodate UTP binding^5^.

Humans have two CTPS isoforms encoded on separate genes, CTPS1 and CTPS2, that share 75% identity. Their relative roles remain unclear. CTPS1 plays a specific and central role in lymphocyte proliferation, and its loss in humans causes severe immune deficiency^15,16^. CTPS is frequently misregulated in cancer^7,17^, with CTPS2 misregulation specifically implicated in osteosarcoma^18^. Given these roles in health and disease, how the two human enzymes are differentially regulated remains an open question of clinical significance.

Polymerization provides an additional layer of CTPS regulation, although the mechanisms by which filaments modulate enzyme activity vary among species. In *E. coli* CTPS filaments stabilize a product-bound, inactive conformation of the enzyme, leading to enhanced inhibition in the filament^1,5^. By contrast, human CTPS1 forms hyper-active filaments composed of enzyme in an active, substrate-bound conformation that disassemble on CTP binding^5^. CTPS filaments appear in response to cellular stress, during particular developmental stages, and in tumor tissue, suggesting a role in adaptation to changing metabolic needs^19–23^.

## RESULTS

### CTPS2 forms distinct substrate- and product-bound filaments

Given the importance of understanding the regulatory differences betwee the two human isoforms and the observed variation in filament-based regulation among species, we aimed to determine whether there are differences in filament structure and function between CTPS1 and CTPS2. We imaged CTPS2 by negative stain EM in the presence of substrates UTP and ATP or products CTP and ADP (Fig. 1a). Surprisingly, unlike CTPS1, which only assembles in the substrate-bound conformation and disassembles upon CTP addition, either substrates or products promoted CTPS2 filament assembly, suggesting a novel mode of regulation.

**Fig. 1:**
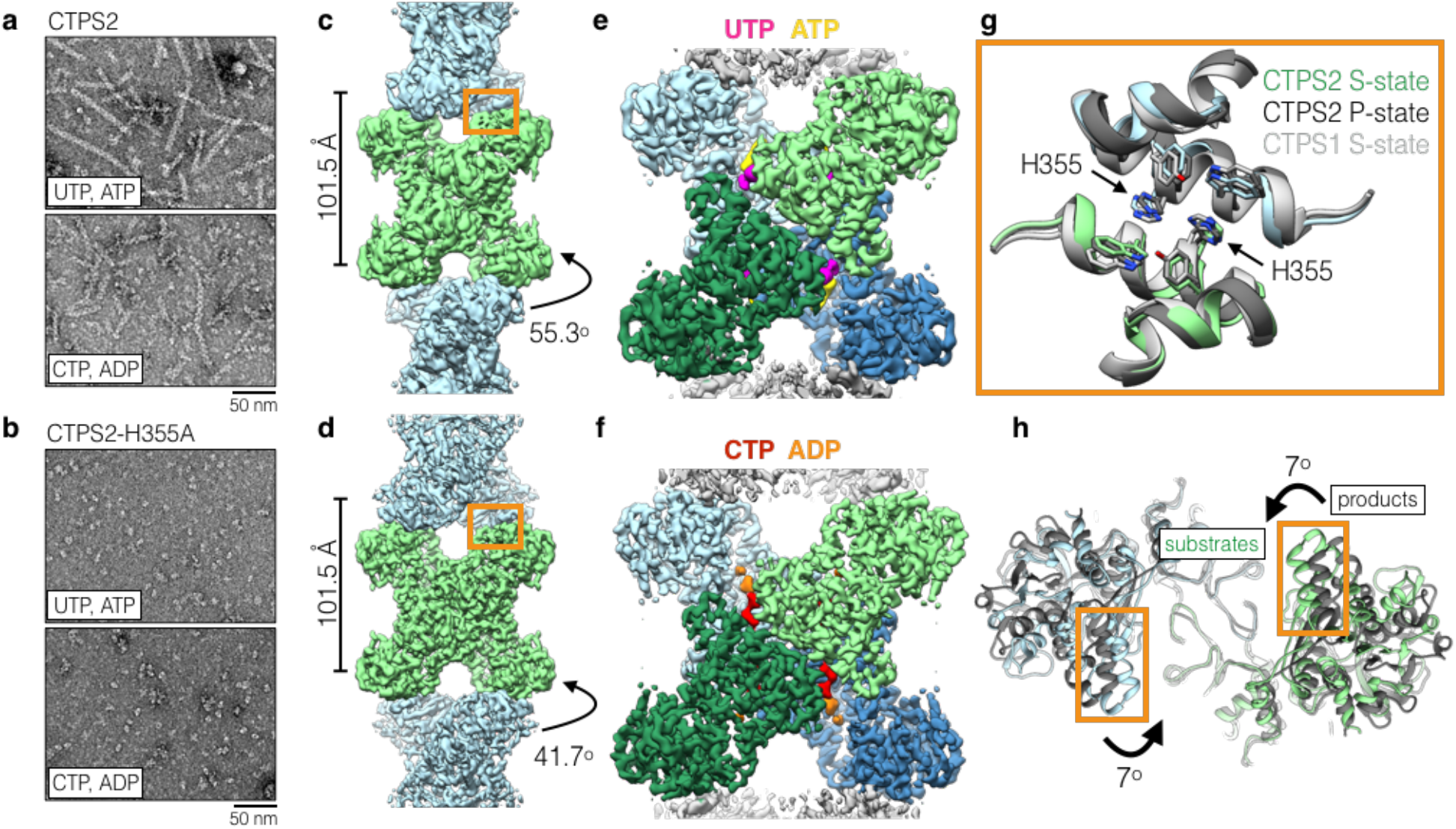
CTPS2 forms distinct S and P-state filaments using the same filament interface. (a,b) Negative stain EM images of CTPS2 wild-type (a) or CTPS2-H355A non-polymerizing mutant (b) in the presence of substrates or products. (c,d) Initial cryoEM reconstructions of S-state (c) (4Å resolution) and P-state (d) (3.6 Å resolution) CTPS2 filaments showing differing helical symmetry, colored by tetramer. Helical rise and rotation are indicated. (e,f) High-resolution reconstructions of S-state (e) (3.5Å resolution) and P-state (f) (3.1 Å resolution) CTPS2 filaments focused on a single tetramer, colored by protomer. Nucleotides are colored as indicated. (g) The filament interface (orange box in c,d) is identical in the CTPS2 S-state (color), CTPS2 P-state (dark grey), and CTPS1 S-state (light grey) structures. Position of conserved H355 is indicated. (h) View down the helical axis comparing the positions of the filament contacts (orange box) in the S-state (color) and P-state (grey) filaments.

We solved cryoEM structures of substrate-bound (S-state) and product-bound (P-state) CTPS2 filaments. Initial reconstructions encompassing multiple tetramers revealed the helical architecture of the filaments (Fig. 1c, d), while masked refinements focused on single tetramers produced higher resolution structures at 3.5Å and 3.1Å of the S-state and P-state filaments, respectively (Fig. 1e, f, Supplementary Fig. 1a-d, Supplementary Table 1). Both CTPS2 filaments are composed of stacked tetramers, with a eukaryote-specific helical insert in the glutaminase domain forming the interfaces between tetramers. The filament assembly interactions are identical in both CTPS2 filament states (Cα RMSD 0.8 Å), and are the same as the CTPS1 interface^5^ (Cα RMSD 1.3 Å) (Fig. 1g). We previously showed that mutation of conserved H355 at the filament interface completely abolishes CTPS1 polymerization^5^. This mutation has the same effect on both S- and P-state CTPS2 polymerization (Fig. 1b). However, CTPS1 filaments have additional interactions between poorly ordered C-terminal tails of adjacent tetramers^5^, which we did not observe in either CTPS2 filament (Supplementary Fig. 1i).

While the filament assembly interfaces are identical in both CTPS2 filament states, the conformation of the enzyme and the helical symmetry are strikingly different. The S-state CTPS2 and CTPS1 filaments are very similar at the level of monomer, tetramer, and filament^5^ (Fig. 1c,e, Supplementary Fig. 1e-h). By contrast, tetramers in the CTPS2 P-state filament are in an inactive, CTP-inhibited conformation, similar to that observed in bacterial CTPS homologs^5,10,14^ (Fig. 1d,f, Supplementary Fig. 1e-h). These differences in conformation result in different helical architectures. The two domains of each protomer rotate relative to each other by 7° between the S- and P-states; the interactions at the interdomain interface remain fixed (Cα RMSD 0.8 Å), with the rotation arising from flexing of residues 40-87 relative to the core of the amidoligase domain (Supplementary Fig. 2a-d). The interdomain rotation alters the positions of filament contacts around the helical axis, leading to a 14^°^ difference in the helical rotation per tetramer between the S-state and P-state filaments (Fig. 1c,d,h). Rare CTPS2 filaments observed in the absence of nucleotides had S-state architecture in negative stain reconstructions, suggesting this may be a somewhat more stable conformation of the enzyme (Supplementary Fig. 3). CTPS2 therefore assembles into active and inactive filaments with unique architectures, depending on the ligand-binding and conformational state of constituent tetramers, while maintaining a fixed filament interface.

### Novel product binding in the P-state CTPS2 filament

The P-state CTPS2 filament structure revealed a novel ADP binding site. In all existing crystal and cryoEM structures of CTPS homologs with adenine nucleotides bound, the adenine ring binds to a pocket formed by R211 and the “lid” residues 238-244^5,10^. In S-state CTPS2 filaments ATP is bound in the same position as in previous structures. By contrast, in the P-state CTPS2 filament, while the ADP phosphates are bound in the conventional position, the adenine base is reoriented by approximately 90° towards the glutaminase domain, and packs in a new site between residues N73 and F77 (Fig. 2a-c). This suggests that the adenine base can bind both sites in CTPS2, and switches to the second site upon transition to the P-state. Furthermore, the overlap between the ATP and ADP binding sites could allow ADP to act as an allosteric regulator, similar to the allosteric regulation observed with CTP at the partially overlapping UTP/CTP binding site.

**Fig. 2:**
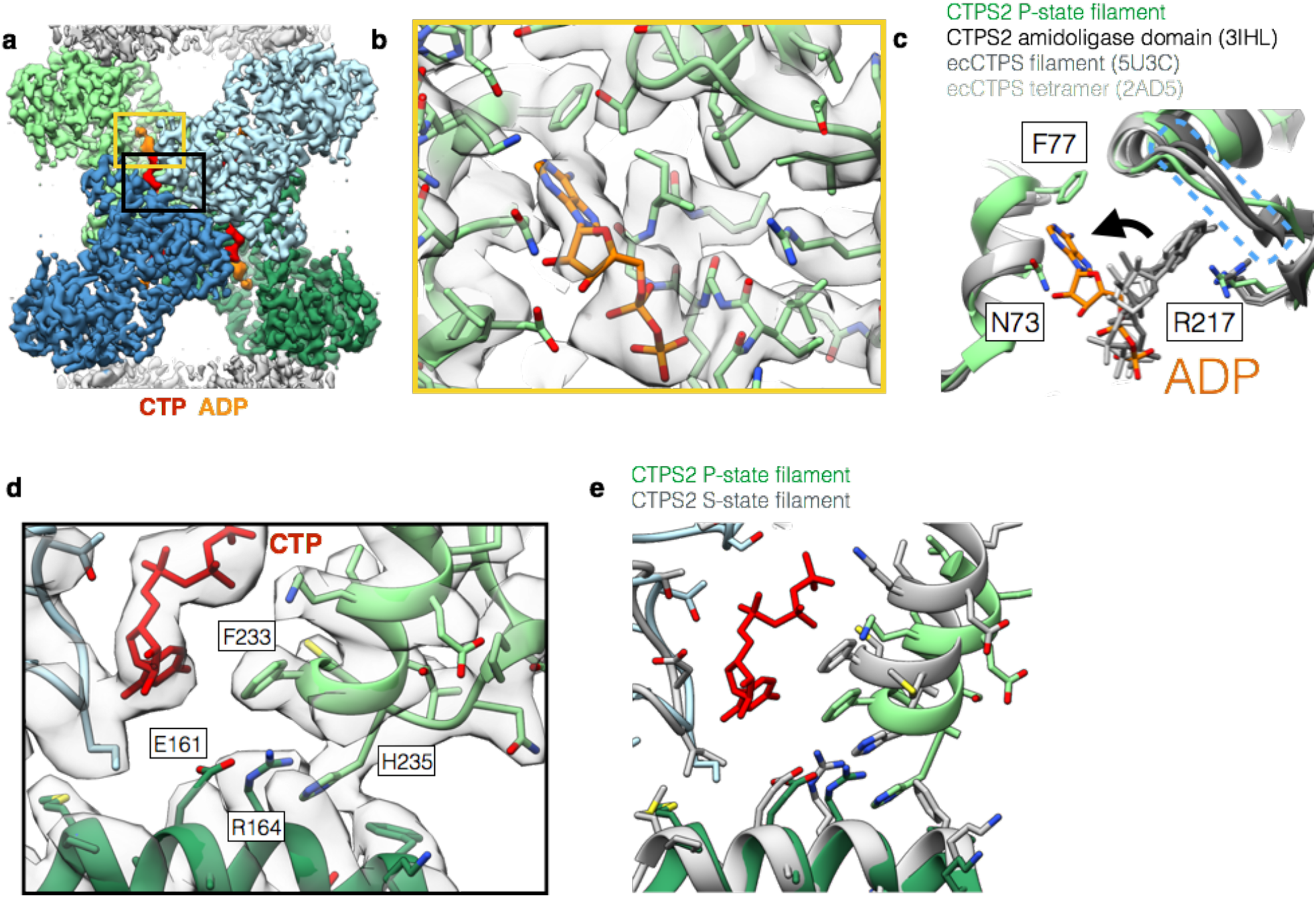
Product binding and changes in the tetramer interface in CTPS2 structures. (a) CTPS2 P-state tetramer, colored by monomer. (b) Zoomed-in view of the yellow box in (a), showing ADP (orange) bound in a novel conformation. cryoEM density is shown in transparent grey. (c) Comparison of ADP conformations in the CTPS2 P-state filament (color) with existing ADP-bound CTPS structures (grey). ADP in the P-state filament is packed between residues F77 and N73. In other CTPS structures, ADP is bound to a pocket formed by R217 and lid residues 244-250 (dashed blue box). (d) Zoomed-in view of the black box in (a), showing the CTP binding site. (e) Comparison of the tetramer interface around the CTP binding site in the P-state (blue and green) and S-state (grey) structures.

The CTP-binding mode in the P-state filament structure is the same as that in existing CTP-bound *E. coli* CTPS structures^5,10^: helix 224-234 is pulled towards CTP, with F233 packing against the CTP base, producing a hydrogen-bonding network amongst residues E161, R164, and H235 at the tetramerization interface (Fig. 2d, e). Consistent with this binding mode, mutation of these residues has been shown to eliminate feedback inhibition of CTPS^1^, including of CTPS1 in CHO cells that results in resistance to chemotherapeutic drugs^7^.

### Substrate binding opens a tunnel in the CTPS monomer

The S-state CTPS2 filament structure is the first near-atomic resolution structure of any CTPS in the substrate-bound conformation, providing insight into the mechanism of ammonia transfer between the two active sites. Previous studies have identified a putative ~25 Å tunnel required to facilitate ammonia transfer between the glutaminase and amidoligase active sites^14,24^ (Fig. 3). However, in the P-state CTPS2 structure as well as existing P-state bacterial structures, this tunnel is blocked by a constriction formed by conserved residues V58, P52, and H55 (V60, P54, and H57 in *E. coli*)^14^ (Fig. 3a, c, Supplementary Fig. 4a, c). Based on a crystal structure of *E. coli* CTPS, Endrizzi *et al.^14^* predicted that H57 may act as a “gate” at the exit of the ammonia tunnel, with UTP binding altering the orientation of H57, causing the gate to open. Indeed, in the S-state CTPS2 filament, H55 reorients to interact with the UTP base, pulling loop P52-V58 towards the amidoligase active site (Fig. 3b, d, Supplementary Fig. 4b, d) (Supplementary Video 1). This conformational change opens the H55 gate and relieves the P52-V58 constriction, providing a tunnel with a nearly uniform ~4 Å diameter for ammonia transfer between the two active sites (Fig. 3e-h). This structural coupling of substrate binding with opening of the ammonia tunnel likely provides the mechanistic basis for the observed coupling of the two enzymatic activities of CTPS, which ensures ammonia is only released into the active site when a UTP substrate is present to accept it^25^.

**Fig. 3:**
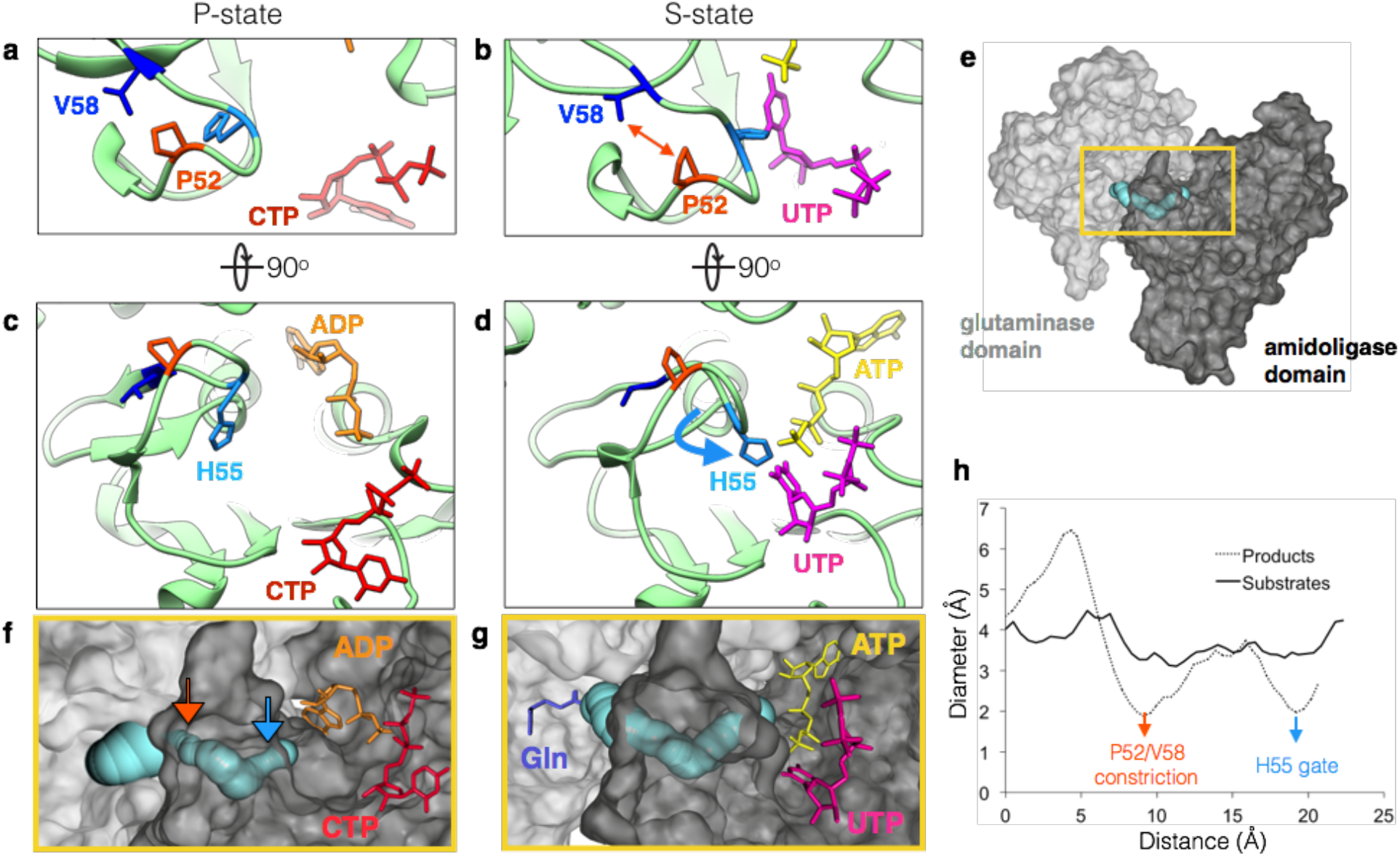
UTP binding opens a tunnel connecting the amidoligase and glutaminase active sites in the S-state CTPS2 filament. (a,b) Conformation of residues at the P52-V58 constriction in the P-state (a) and S-state (b) CTPS2 filaments. (c,d) Position of the H55 gate in the P-state (c) and S-state (d) CTPS2 filaments. (e) CTPS2 monomer showing the position of the tunnel (blue) linking the Glutaminase (light grey) and Amidoligase (dark grey) domains. (f,g) Zoomed-in view of the yellow box in (e), showing tunnels identified by CAVER software in the P-state (f) and S-state (g) CTPS2 structures. Positions of the H55 gate (blue arrow) and P52-V58 constriction (orange arrow) in the P-state structure are indicated. The expected position of glutamine is shown in purple (based on PDB 1VCO). (h) Plot showing the diameter of the tunnel along its length for P and S-state structures, with positions of the H55 gate (blue arrow) and P52-V58 constriction (orange arrow) indicated.

### Regulation of CTPS2 filaments is highly cooperative

Given that the filament interface is identical in the S-state and P-state structures, we hypothesized that CTPS2 filaments could directly switch between the S-state and P-states while remaining polymerized, perhaps allowing for coordinated conformational changes along entire filaments. To test this hypothesis, we trapped CTPS2 in filaments by engineering cysteine disulfide crosslinks at the filament interface, yielding the CTPS2^CC^ mutant (Fig. 4a, b, Supplementary Fig. 5). In the absence of ligands CTPS2^CC^ spontaneously and robustly polymerized into filaments under non-reducing conditions (Fig. 4c). We generated 2D averages of crosslinked CTPS2^CC^ in different ligand states to probe for conformational changes within the filaments. Because their different helical symmetries give rise to a characteristic ~180° repeated view every 300 Å (S-state) or 400 Å (P-state), the different architectures are readily distinguishable in 2D averages (Fig. 4d). Classification and alignment of 2D averages to low-pass filtered projections of the CTPS2 structures revealed that apo CTPS2^CC^ filaments had S-state architecture and transitioned to the P-state upon addition of CTP, confirming that conformational switching within intact filaments is possible (Fig. 4e) (Supplementary Video 2).

**Fig. 4:**
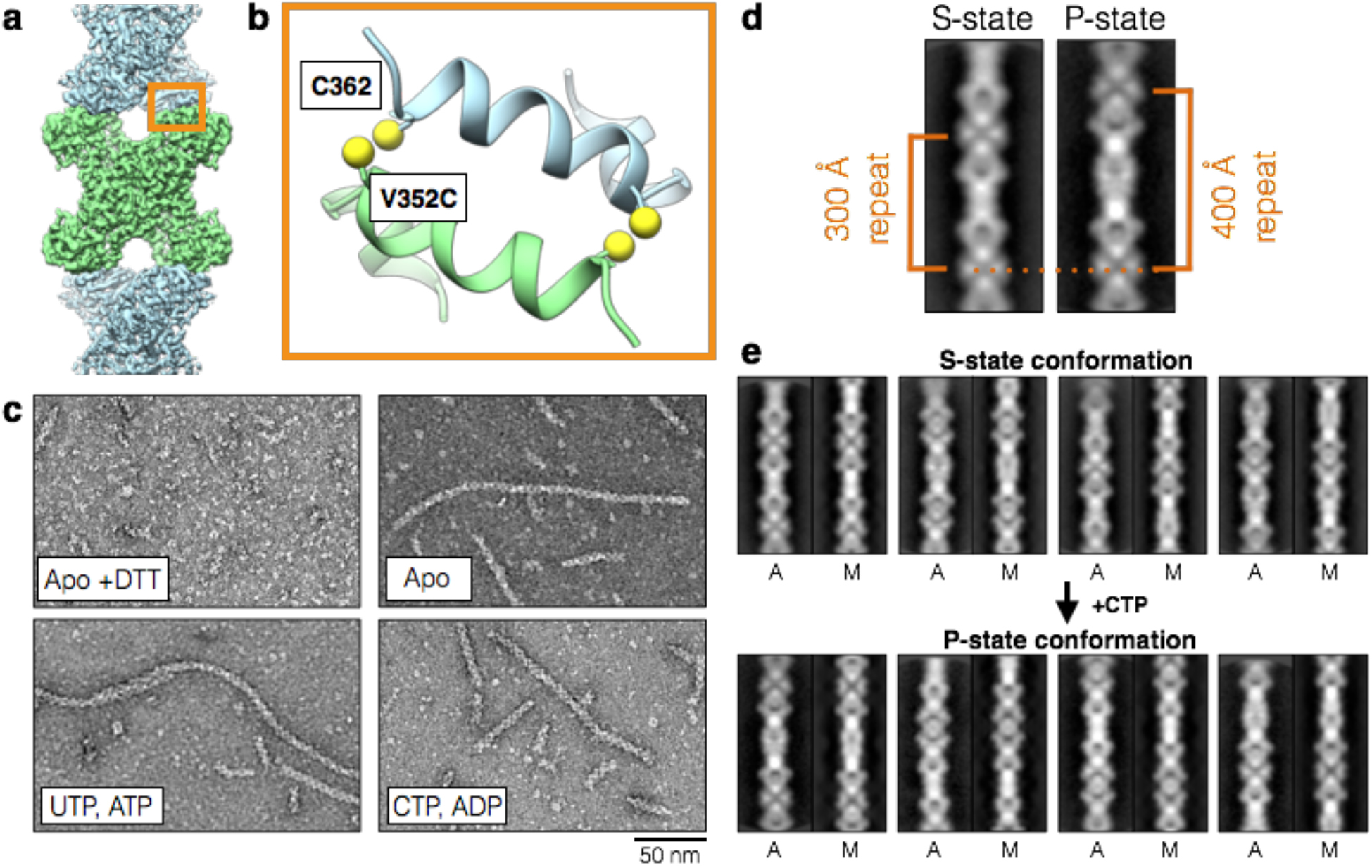
Disulfide-crosslinked CTPS2 filaments can switch conformations. (a) CTPS2 filament showing the position of the CTPS2^CC^ mutant at the filament interface. (b) Zoomed-in view of the orange box in (a), showing the design of the CTPS2^CC^ mutant. The V352C mutation is across from native C362. (c) Negative stain EM images of hCTPS^CC^ under reducing conditions (+DTT), and under non-reducing conditions in the presence or absence of nucleotides. (d) 2D averages of S and P-state CTPS2^CC^ filaments, with characteristic 180° repeat distances indicated. (e) 2D class averages of cross-linked CTPS2^CC^ filaments before and after addition of CTP. 2D averages (A) are aligned to low-pass filtered projections of CTPS2 cryoEM models (M) in the S-state (top) and P-state (bottom) conformations.

We suspected that linking the conformational state of many CTPS2 subunits within a filament could lead to enhanced cooperativity in CTPS2 regulation. We therefore compared the UTP and CTP kinetic parameters of CTPS2 filaments with those of the CTPS2-H355A non-polymerizing mutant (Fig. 5, Supplementary Fig. 6). Unlike CTPS1^5^, polymerization did not increase the V_max_ of CTPS2 (Fig. 5a). Further, CTPS2 filaments and CTPS2-H355A homotetramers exhibited nearly identical S_0.5_ and IC_50_ values for UTP and CTP (Fig. 5b-d, Supplementary Table 2). CTPS2-H355A inhibition is highly cooperative, with n_Hill_ of 3.5 that approaches the theoretical limit for a tetramer (Fig. 5c). CTPS kinetic parameters reported for various species vary significantly, but hill coefficients close to 4 have been reported for both activation and inhibition^26,27^. Remarkably, CTPS2 filaments exhibited an even higher n_Hill_ of 8.3, providing a switch-like response to changes in CTP concentrations (Fig. 5c). Dilution of wild-type CTPS2 below its critical concentration for assembly caused disassembly into tetramers and occasional short filaments, resulting in a n_Hill_ of 3.9, similar to that observed for the non-polymerizing mutant (Fig. 5d, f). Polymerization therefore greatly increases the cooperativity of CTPS2 regulation, likely due to concerted conformational changes within filaments (Fig. 5h).

**Fig. 5:**
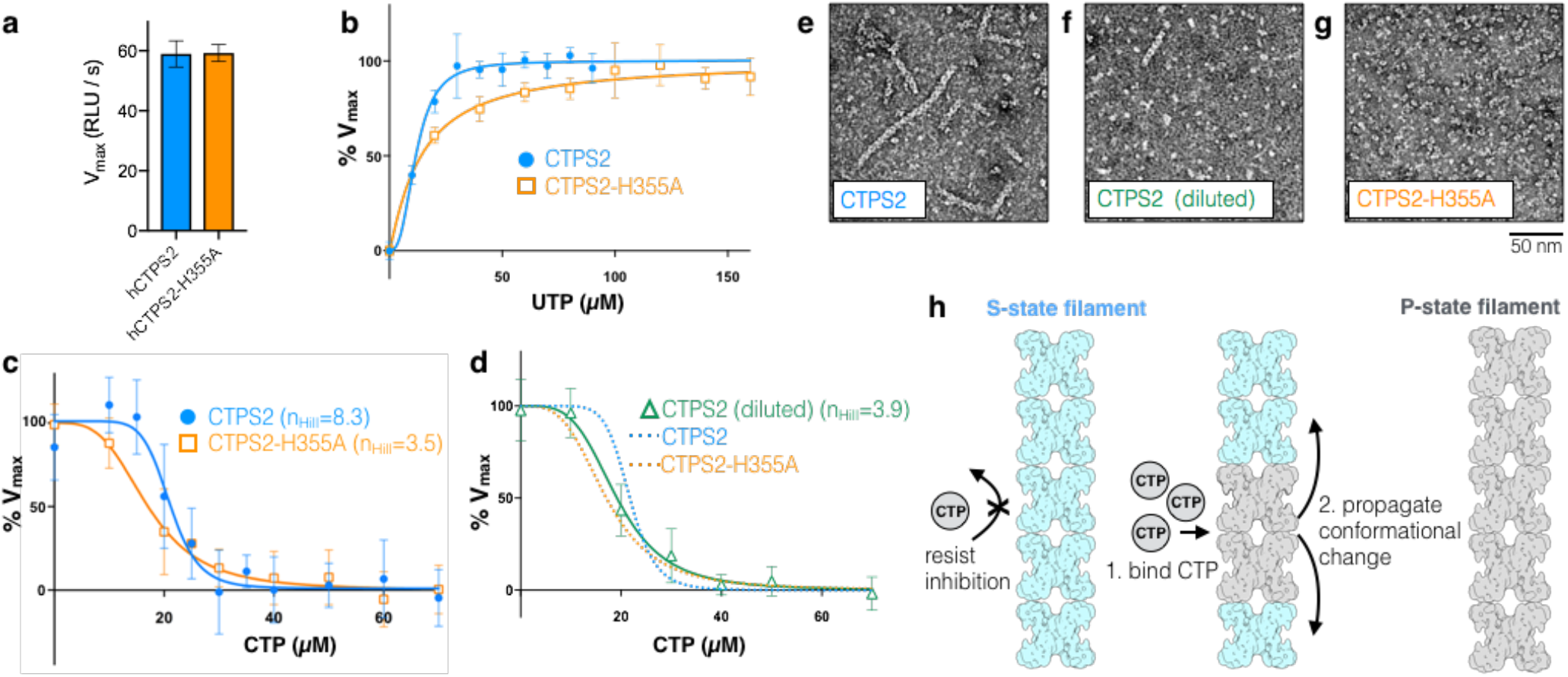
Regulation of CTPS2 filaments is highly cooperative. (a-d) ADP-Glo enzyme assays comparing kinetics of CTPS2 filaments and tetramers. (a) V_max_ is the same for CTPS2 filaments and CTPS2-H355A tetramers. (b) UTP kinetic curves for CTPS2 and CTPS2-H355A. Assays were performed with 1500 nM enzyme. (c) CTP inhibition curves for CTPS2 and CTPS2-H355A at 1500 nM protein. n_Hill_ values are indicated. (d) CTP inhibition curve for CTPS2 at 300 nM protein. Fits for CTPS2 and CTPS2-H355A inhibition from panel (c) are reproduced for comparison. n_Hill_ value is indicated. (e-g) Negative stain EM images of CTPS2 and CTPS2-H355A under ADP-Glo assay conditions. CTPS2 forms filaments at 1500 nM concentration (e), but disassembles into tetramers and rare short filaments at 300 nM concentration (f). CTPS2-H355A does not form filaments at 1500 nM concentration (g). (h) Model for cooperative regulation in hCTPS2 filaments. CTP-binding induces conformational changes which propagate along filaments. Assays were performed in triplicate (6 assays for CTPS2 in (c)) and measured in triplicate, and values are presented as mean +/- s.d.

## DISCUSSION

Given that the residues involved in filament assembly interactions are completely conserved between CTPS1 and CTPS2, it is intriguing that CTPS1 forms S-state but not P-state filaments^5^. It is possible that product-bound CTPS1 adopts a unique conformation incompatible with filament assembly. Alternatively, the contact between disordered C-terminal regions observed in CTPS1^5^ but not CTPS2 may play a role in differential assembly regulation; the C-terminus regulates CTPS2 activity in a phosphorylation-dependent manner and accounts for much of the sequence variation between isoforms^28,29^. Differences in the regulatory role of filaments between CTPS1 and CTPS2 may reflect their different physiological roles: consistent with its role in cellular proliferation during the immune response CTPS1 may be induced into filaments which reduce sensitivity to feedback inhibition and allow expansion of CTP pools to meet increased demand, whereas under more homeostatic conditions the extreme sensitivity of CTPS2 fiilaments may help maintain a strictly defined UTP/CTP balance.

Cooperativity in biological systems often arises from the association of protein subunits into oligomeric complexes, allowing for coordinated regulation through coupling of conformational states. Typically, cooperativity is associated with assemblies of relatively few protein subunits, with hemoglobin tetramers providing a canonical example. Large protein arrays and polymers hold the potential for massive cooperativity, with conformational information integrated across hundreds or thousands of protein subunits^30–32^. Several examples of this phenomenon occur in membrane-embedded systems where two-dimensional arrays can exhibit switchlike transitions, like the chemostatic network controlling bacterial flagellar motion where conformational states propagate across clusters of membrane-bound receptors to amplify external signals^33–35^, or the coupled gating of ryanodine receptor arrays in muscle cells^36^. In the case of CTPS2, highly cooperative enzyme regulation results from ligand-induced propagation of conformational changes along linear polymers. This allows for increased sensitivity to substrate and product balance. Sensitivity in the regulation of nucleotide biosynthesis is important to the maintenance of genomic integrity as imbalances in nucleotide pools are linked to increased mutagenesis, sensitization to DNA damaging agents, and multidrug resistance^37,38^.

Many enzymes in a broad range of metabolic pathways form filamentous polymers^19,21,39-42^. In the few examples that have been biochemically and structurally characterized to date, enzyme regulation arises from assembling filaments that stabilize particlular conformational states^1–5^. The enhanced cooperativity of CTPS2 is a novel filament-based mechanism of enzyme regulation, which likely serves to stabilize nucleotide levels over a narrow concentration range. This new function for metabolic filaments highlights the diversity of ways in which self-assembly can be adapted to allosterically fine-tune enzyme regulation and improve the efficiency of metabolic control.

## Acknowledgments

We thank G. Carman (Rutgers University) for the hCTPS2-expressing *S. cerevisiae* strain. We are grateful to the Arnold and Mabel Beckman Cryo-EM Center at the University of Washington for use of electron microscopes.

## Data availability

EM structures and atomic models have been deposited in the Electron Microscopy Data Bank and Protein Data Bank with the accession codes: S state CTPS2 filament (EMD-20354; PDB 6PK4), P state CTPS2 filament (EMD-20355; PDB 6PK7).

## Author contributions

E.M.L. performed the experiments. E.M.L. and J.M.K designed experiments, performed data analysis and interpretation, and wrote the manuscript.

## METHODS

### Purification of CTPS2

CTPS2 was expressed in *Saccharomyces cerevisiae* strain GHY56, as described by Han *et al^43^*. In GHY56, both endogenous *Saccharomyces cerevisiae* CTP synthase genes URA7 and URA8 are deleted. This lethal deletion is rescued by plasmid pDO105-CTPS2, which directs expression of C-terminally His_6_-tagged CTPS2 from the ADH1 promoter. GHY56 cells were grown in 4X YPD at 30°C, harvested by centrifugation, frozen as droplets in liquid nitrogen, and then ground into powder while frozen. Cell powder (~20g) was resuspended in lysis buffer (50 mM Tris-HCl, 200 mM NaCl, 0.3M sucrose, 20 mM imidazole, pH 8.0), and centrifuged at 14,000 RPM for 40 minutes at 4°C in a Thermo Scientific Fiberlite F14-14 × 50cy rotor. Clarified lysate was loaded onto a 5 mL HisTrap FF Crude column (GE) on an ÄKTA Start chromatography system (GE) pre-equilibrated in column buffer (20 mM Tris-HCl, 0.5M NaCl, 45 mM imidazole, 10% glycerol, pH 7.9). The column was washed with 25 column volumes (CV) of column buffer, and CTPS2 was eluted with 5CV of elution buffer (20 mM Tris-HCl, 0.5M NaCl, 250 mM imidazole, 10% glycerol, pH 7.9) as 1 mL fractions. Fractions containing CTPS2 were pooled and dialyzed into storage buffer (20 mM Tris-HCl, 0.5M NaCl, 10% glycerol, 7 mM β-mercaptoethanol, pH 7.9) using Snakeskin 3500 MWCO dialysis tubing (Thermo Scientific). Dialyzed CTPS2 was then concentration approximately 5-fold using a 3 kDa cut-off centrifugal filter unit (Millipore), flash-frozen in liquid nitrogen, and stored at −80°C. CTPS2 mutants were purified using the same methods.

### Cloning of mutants

CTPS2-H355A and CTPS2^CC^ mutants were generated using the Gibson assembly method^44^. PCR was used to amplify two separate fragments of the pDO105-CTPS2 plasmid backbone flanking the mutation site, which were then ligated together with ~60 base pair DNA fragments containing the desired H355A or V352C mutations. Mutations were confirmed by Sanger sequencing.

### Negative-stain electron microscopy

Prior to imaging, CTPS2 in storage buffer was exchanged into imaging buffer (20 mM Tris-HCl, 100 mM NaCl, 7 mM β-mercaptoethanol, pH 7.9) using a 7K MWCO Zeba Spin Desalting Column (Thermo Scientific). For imaging CTPS2^CC^ disulfide crosslinked filaments, protein was exchanged into non-reducing buffer (20 mM Tris-HCl, 100 mM NaCl, pH 7.9). 100 mM DTT was added to depolymerize CTPS2^CC^ filaments. CTPS2 was applied to glow-discharged carbon-coated grids, stained with 0.7% uranyl formate, and imaged on a Tecnai G2 Spirit (FEI co.) operating at 120 kV. Images were acquired at 67,000× magnification on a US4000 4k × 4k CCD camera (Gatan, Inc.).

### Negative-stain electron microscopy image processing and reconstructions

3D reconstruction of Apo CTPS2 filaments was performed using iterative helical real space reconstruction (IHRSR)^45,46^ in SPIDER, with hsearch_lorentz^47^ used to refine helical symmetry parameters, and with D2 point-group symmetry enforced. Cryo-EM structures of S-state or P-state CTPS2 filaments low-pass filtered to 40Å were used as starting models. 2D class averages of CTPS2^CC^ were generated by manually picking particles and performing 2D classification in Relion^48^. CTPS2^CC^ class averages were aligned to 30Å low-pass filtered projections of the S-state and P-state cryoEM structures using e2classvsproj.py in EMAN2^49^.

### Cryo-electron microscopy

Cryo-EM samples were prepared by applying CTPS2 to glow-discharged CFLAT 1.2/1.3 holey-carbon grids (Protochips Inc.), blotting with a Vitrobot (FEI co.), and plunging into liquid ethane. CTPS2 was exchanged into imaging buffer and incubated with nucleotides for 5 minutes at 37°C before preparing cryo-EM samples. Conditions for the S-state filament structure were 7 µM CTPS2, 2 mM UTP, 2 mM ATP, 0.2 mM GTP, and 10 mM MgCl_2_. Conditions for the P-state filament structure were 8 µM CTPS2, 2 mM CTP, 2 mM ADP, and 10 mM MgCl_2_. Data for preliminary 3D reconstructions was collected on a Tecnai G2 F20 (FEI co.) operating at 200 kV. Movies were acquired on a K-2 Summit Direct Detect camera in counting mode with a pixel size of 1.26 Å/pixel, collecting 36 frames with a total dose of 45 electrons/Å^2^, with a defocus range of −1.0 to −2.5 µm. Data for high-resolution structures was collected on a Titan Krios (FEI co.) equiped with a Quantum GIF energy filter (Gatan Inc.) operating in zero-loss mode with a 20 eV slit width. Movies were acquired on a K-2 Summit Direct Detect camera in super-resolution mode with a pixel size of 0.525 Å/pixel, collecting 50 frames with a total dose of 90 electrons/Å^2^. Movies were collected within a defocus range of −1.0 to −2.5 µm. EPU (FEI co.) and Leginon^50^ software were used for automated data collection.

### Cryo-electron microscopy image processing and reconstructions

Movie frame alignment and dose-weighting was performed using MotionCor2^51^, and CTF parameters were estimated using GCTF^52^. For data collected on the Tecnai G2 F20, particles were picked manually using Appion^53^. 3D reconstructions at ~8Å resolution were generated using IHRSR^45,46^ in SPIDER, using cylinders as starting models and imposing D2 symmetry, with helical symmetry refined using hsearch_lorentz^47^. For Titan Krios data, particles were initially picked manually from a subset of images, and used to generate 2D averages in Relion^48^. These initial 2D averages were used as templates for Relion automated picking from all images. 2D classification in Relion was used to remove poorliy aligning particle picks, and well-defined particles were then subject to Relion auto-refinement, using the SPIDER reconstructions low-pass filtered to 30Å as starting models. D2 symmetry was imposed during all 3D refinement. Parameters, particles, and structures from Relion auto-refine were exported to cisTEM^54^, where multiple further rounds of 2D and 3D classification were performed. Final reconstructions of S-state and P-state CTPS2 filaments were generated in cisTEM, using automatic refinement, followed by manual refinement with CTF refinement implemented. Maps were sharpened in cisTEM using a b-factor of 50 Å^2^. Resolutions were estimated using the FSC_0.143_ cutoff. Details of 3D reconstructions are summarized in Supplementary Table 1.

### Model building

MODELLER^55^ was used to generate an initial homology model of the full-length CTPS2 monomer, using partial crystal structures of the CTPS2 glutaminase (PDB 2V4U) and amidoligase domains (PDB 2VO1) aligned to a crystal structure of the full-length *E. Coli* CTPS monomer (PDB 2AD5). The CTPS2 glutaminase, amidoligase, and linker domains were fit individually as rigid bodies into EM maps using Chimera. Structures were then refined using multiple cycles of real-space refinement in Phenix^56^ and Coot^57^.

### Tunnel modelling

CAVER Analyst^58^ software was used to model tunnels through the CTPS2 atomic models. The same starting coordinates were used for the S-state and P-state filaments, at a site adjacent to the glutaminase domain catalytic cysteine 399. Probe radii of 0.5Å and 1.5Å were used for the P and S-state structures, respectively, and other tunnel computation parameters were set to default values. The use of a less restrictive, smaller probe radius for the inhibited state allowed us to define a continuous tunnel through the constriction points. Plots of tunnel diameter versus distance were also produced in CAVER Analyst.

### CTPS2 kinetics

Kinetic parameters for CTPS2 and CTPS2-H355A were determined using the ADP-Glo assay (Promega), using similar conditions to those described by Sakamoto *et al^59^*. Assays were performed in 50 mM K-HEPES, 5 mM KCl, 0.01% tween 20, 0.01% BSA, 20 mM MgCl_2_, pH 8.0. All steps of the assay were performed at room temperature in black, low volume 384 well plates (Corning). UTP kinetic assays were performed with 1500 nM CTPS2, 0-150 µM UTP, 500 µM ATP, 5 µM GTP, and 500 µM Glutamine. CTP inhibition assays were performed with 300 nM or 1500 nM CTPS2, 0-70 µM CTP, 100 µM UTP, 100 µM ATP, 5 µM GTP, and 100 µM Glutamine. The total volume of the CTPS2 assays was 6 µl, and reactions were ran for 60 minutes (300 nM CTPS2) or 12 minutes (1500 nM CTPS2). CTPS2 reactions were terminated by addition of 6 µl ADP-Glo reagent and incubated for 1 hour, after which 12 µl of kinase detection reagent was added. After one hour, luminescence was recorded using a Varioskan Lux (Thermo Scientific) microplate reader. Assays were performed in triplicate, and three luminescence readings were taken for each assay and averaged. Kinetics data were fit by 4 parameter logistic regression, solving for maximum rate, minimum rate, hill number, and IC_50_ or S_0.5_. Data were plotted as percent maximum rate, according to the formula: 100*[(observed - minimum)/(maximum - minimum)].

## Video legends

**Supplementary Video 1:** Animation of the CTPS2 active site morphing between the S-state and P-state conformations. The glutaminase domain (green) rotates towards the amidoligase domain (blue), while loop P52-V58 (orange) is pulled towards the UTP base (magenta) in the S-state. ATP (yellow) is also shown in the active site.

**Supplementary Video 2:** Animation of CTPS2 morphing between the S-state and P-state filament conformations. Colored by tetramer. Filament contact sites (orange) remain the same despite conformation changes between the two filament states.

**Supplementary Fig. 1:**
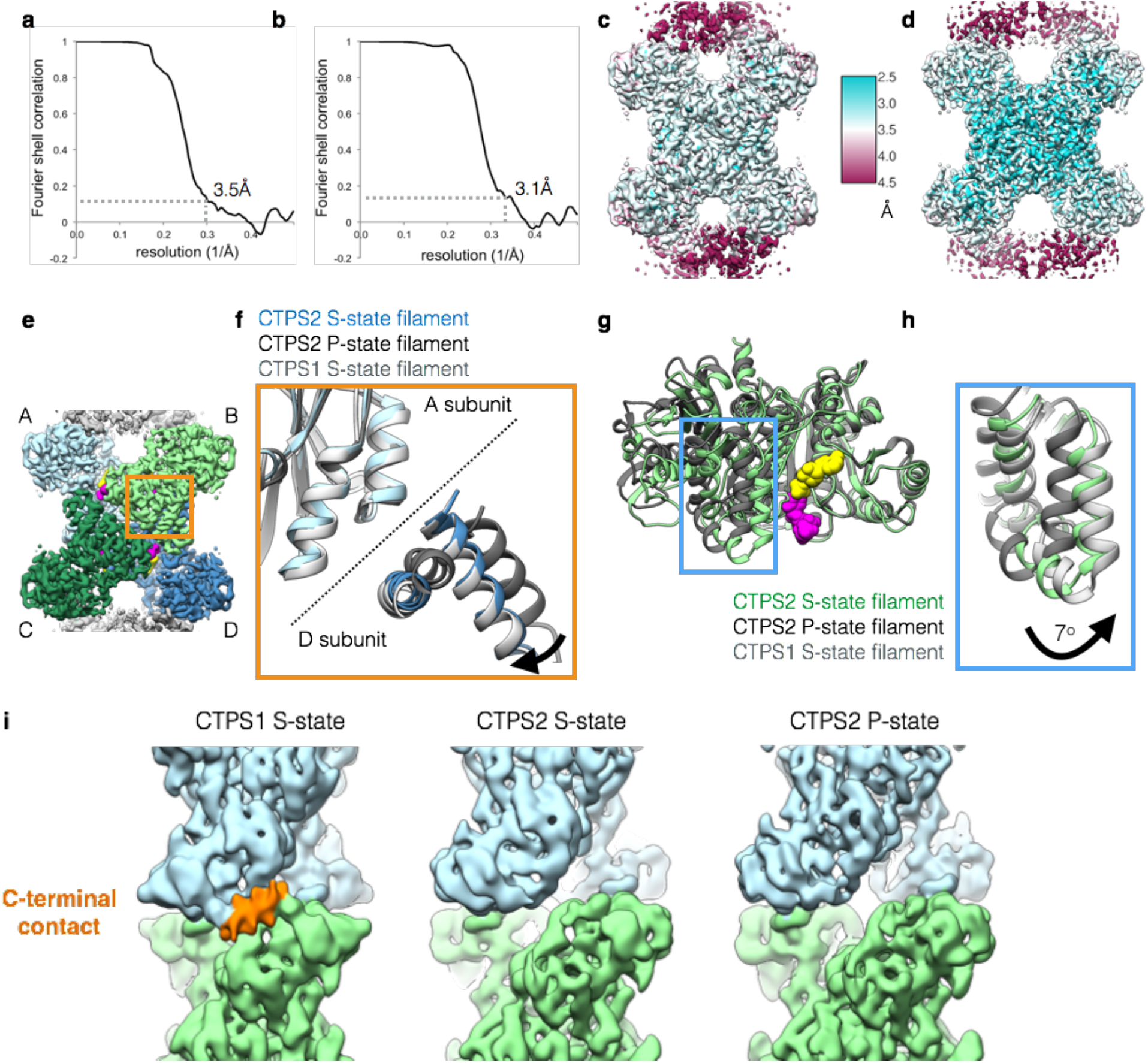
Details of CTPS2 cryoEM structures. (a,b) FSC curves for the S-state (a) and P-state (b) CTPS2 filament structures, showing resolutions of 3.5Å and 3.1Å, respectively, by the FSC_0.143_ criteria. (c,d) ResMap local resolution maps of the S-state (c) and P-state (d) CTPS2 filament structures. (e) CTPS2 tetramer, with monomers A-D shown in different colors. (f) Zoomed-in view of the orange box in (e), showing S-state CTPS2 (blue), P-state CTPS2 (dark grey), and S-state CTPS1 (light grey) filament structures aligned on the Amidoligase domain of subunit A. S-state CTPS1 and CTPS2 are extended across the tetramer interface, compared to P-state CTPS2. (g) Monomers from the S-state (green) and P-state (dark grey) CTPS2 filament structures aligned on the Amidoligase domain. (h) Zoomed-in view of the blue box in (g), with S-State CTPS1 also shown in light grey. The glutaminase domain is rotated by 7° towards the Amidoligase domain in the S-state structure. (i) CryoEM maps of CTPS1 and CTPS2 filaments, low-pass filtered to 8Å for comparison. CTPS1 filaments have an additional, poorly ordered, C-terminal filament contact (orange) that is not observed in S- or P-state CTPS2 filaments.

**Supplementary Fig. 2:**
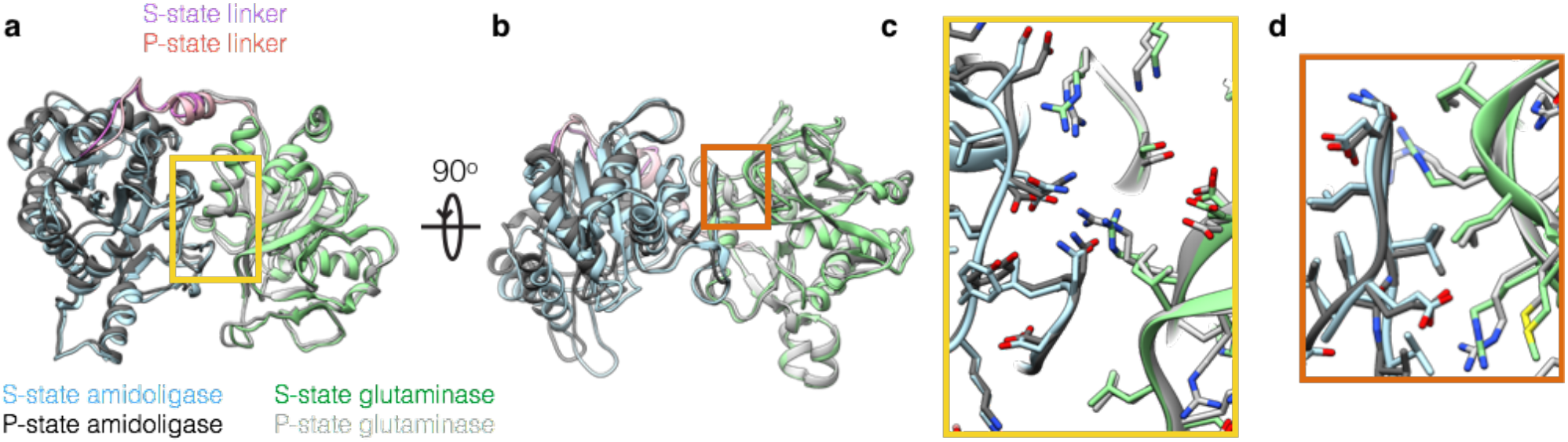
Comparison of the Glutaminase-Amidoligase interface in S- and P-state CTPS2 filaments. (a,b) Two views of the S-state (color) and P-state (grey) CTPS2 monomers aligned on the glutaminase-amidoligase interface. (c) Zoomed-in view of the yellow box in (a). (d) Zoomed-in view of the orange box in (b). The glutaminase-amidoligase interface is essentially identical (Cα RMSD 0.8 Å) in the S-state and P-state filaments.

**Supplementary Fig. 3:**
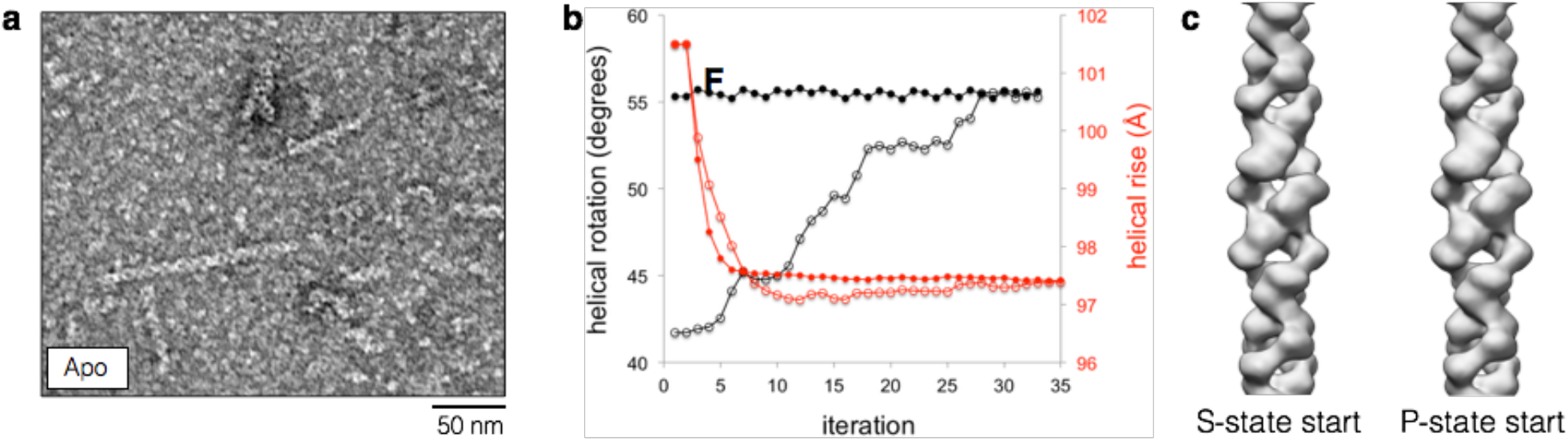
Rare apo CTPS2 filaments have S-state filament architecture. (a) Negative stain EM images of apo CTPS2. Occasional filaments are observed. (b) Helical rise (red) and rotation (black) values plotted over multiple rounds of iterative helical real space reconstruction of apo CTPS2 filaments in stain. Starting helical symmetry values and models from cryoEM structures of S-state (closed circles) or P-state (open circles) CTPS2 filaments were used. Both reconstructions converge on the S-state filament helical symmetry values. (c) The apo CTPS2 filament structures from the reconstructions described in (b) are the same and have the S-state helical rotation, regardless of which starting symmetries and models are used.

**Supplementary Fig. 4:**
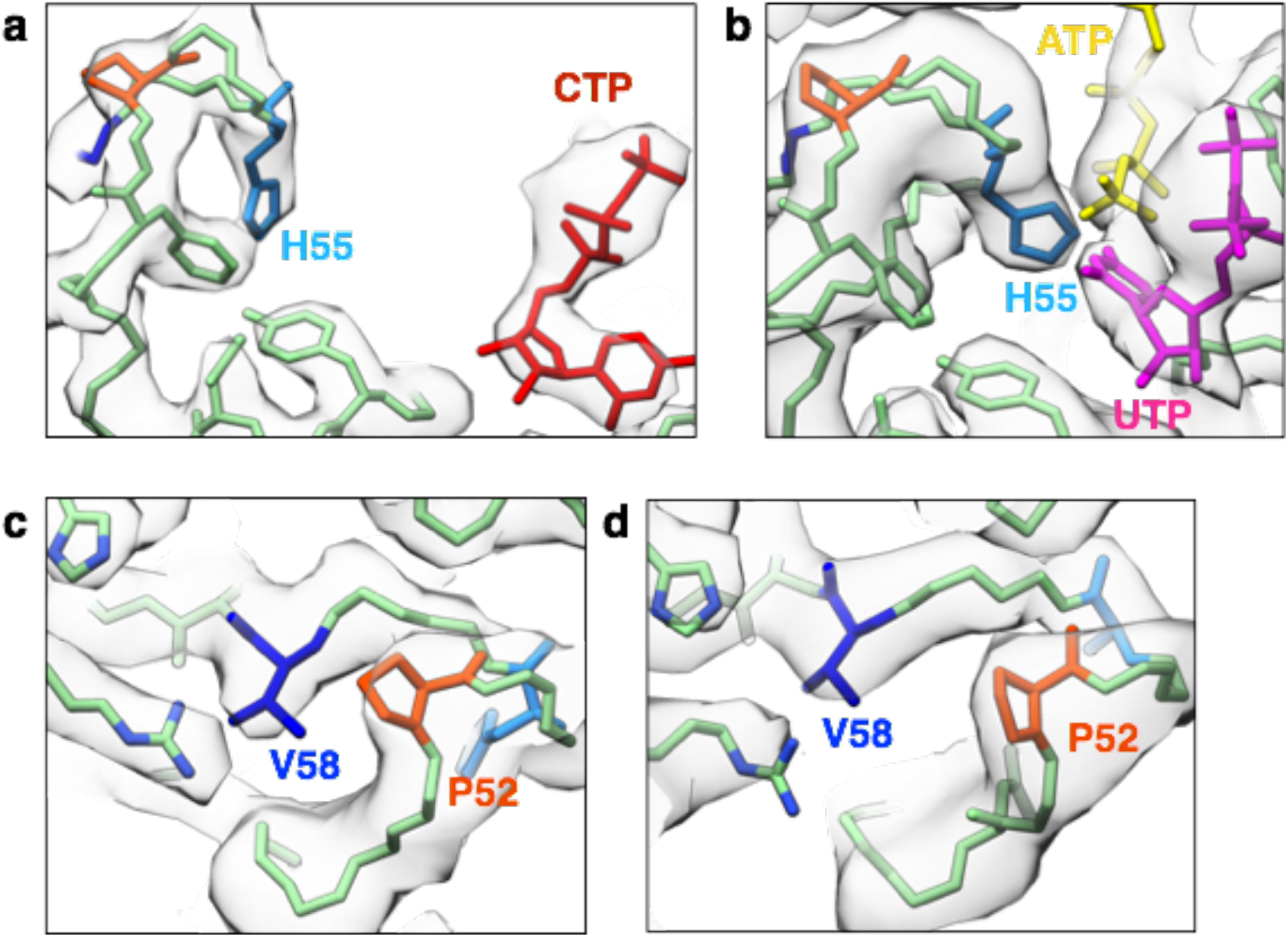
CryoEM density at the P52-V58 and H55 constriction points. (a,b) CryoEM density (grey) and atomic models (color) at the H55 gate in the P-state (a) and S-state (b), CTPS2 filaments. (c,d) CryoEM density (grey) and atomic models (color) at the P52-V58 constriction in the P-state (c) and S-state (d) CTPS2 filaments.

**Supplementary Fig. 5:**
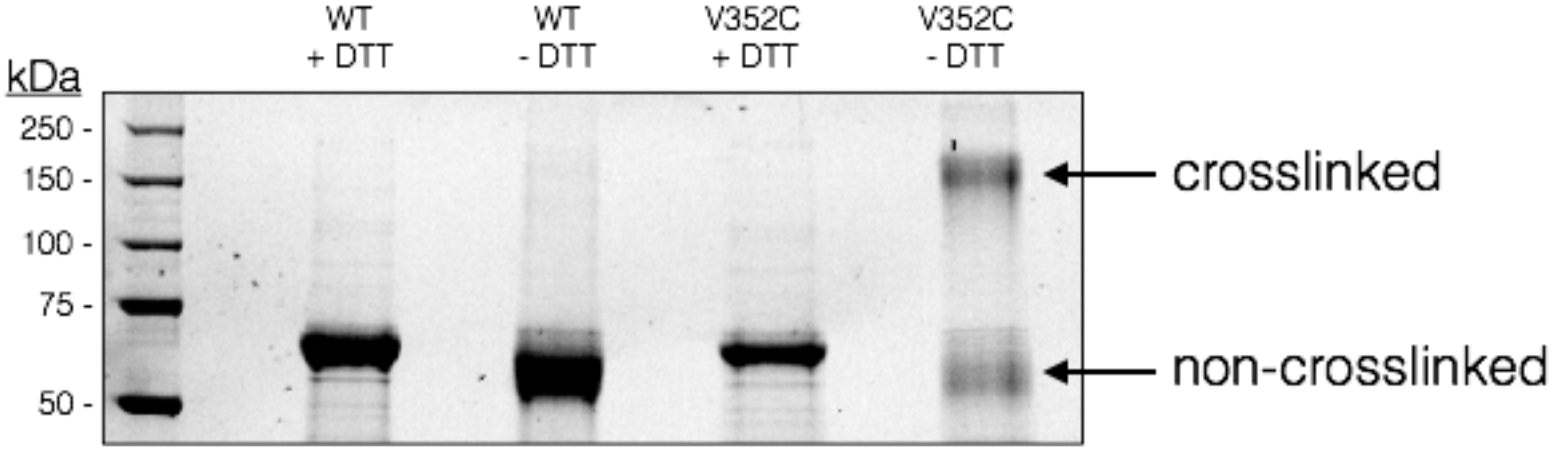
SDS-PAGE gel of wild-type CTPS2 and CTPS2^CC^ under reducing (100 mM DTT) and non-reducing conditions. Bands for crosslinked CTPS2^CC^ are visible under non-reducing conditions.

**Supplementary Fig. 6:**
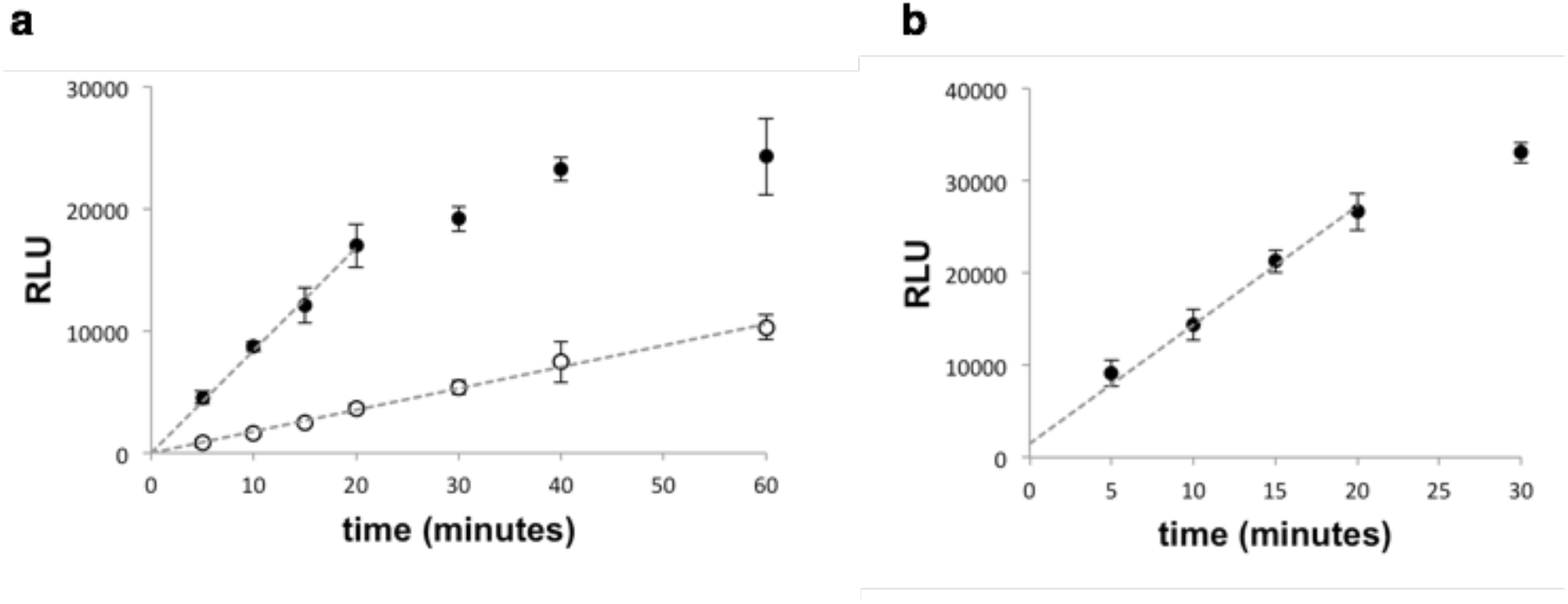
ADP-Glo assays show a linear response in relative light units (RLU) over the time used for CTPS2 kinetics assays. (a) Time-course of ADP-Glo assay under the substrate conditions used for CTP inhibition assays at 300 nM CTPS2 (open circles) and 1500 nM CTPS2 (closed circles). (b) Time-course of ADP-Glo assay at 1500 nM CTPS2 with the lowest UTP concentration used in UTP kinetics assays (10 µM UTP). Dashed lines indicate the linear portion of the assay used to measure reaction velocity. Assays were performed in triplicate and measured in triplicate, and values are presented as mean +/− s.d.

**Supplementary Table 1:** EM data collection and refinement statistics

|  | **S state CTPS2 filament**<br>(EMD-20354, PDB 6PK4) | **P state CTPS2 filament**<br>(EMD-20355, PDB 6PK7) |
| --- | --- | --- |
| **Data collection** |  |  |
| Electron microscope | Titan Krios | Titan Krios |
| Voltage (kV) | 300 | 300 |
| Electron detector | K2 summit | K2 summit |
| Electron dose (e<sup>-</sup>/Å<sup>2</sup>) | 90 | 90 |
| Pixel size (Å) | 1.05 | 1.05 |
| **Reconstruction** |  |  |
| Point group symmetry | D2 | D2 |
| Helical symmetry (rise, rotation) | 101.5 Å, 55.3° | 101.5 Å, 41.7° |
| Particles | 53964 | 22705 |
| Resolution (0.143 fsc) (Å) | 3.5 | 3.1 |
| **Refinement** |  |  |
| Initial model used | PDB 2V4U, PDB 2VO1 | PDB 2V4U, PDB 2VO1 |
| **Model composition** |  |  |
| Protein residues | 559 | 557 |
| Ligands | UTP, ATP, Mg | CTP, ADP, Mg |
| **Validation** |  |  |
| Clashscore | 12 | 11 |
| Poor rotamers (%) | 1 | 0.4 |
| **Ramachandran plot** |  |  |
| Favored (%) | 92 | 91 |
| Allowed (%) | 8 | 9 |
| Outliers (%) | 1 | 0 |

**Supplementary Table 2:** Summary of CTPS2 kinetic parameters

|  | UTP kinetics |  | CTP inhibition |  |
| --- | --- | --- | --- | --- |
|  | S<sub>0.5</sub> (μM) | n<sub>Hill</sub> | IC<sub>50</sub> (μM) | n<sub>Hill</sub> |
| CTPS2 (1500 nM) | 11.7 | 2.8 | 21.0 | 8.3 |
| CTPS2-H355A (1500 nM) | 14.0 | 1.1 | 17.1 | 3.5 |
| CTPS2 (300 nM) | - | - | 19.2 | 3.9 |

